# Fully Two-Photon-Driven RESOLFT Microscopy for Super-Resolution Imaging inside Tissue

**DOI:** 10.64898/2026.09.13.751305

**Authors:** Ryohei Ozaki-Noma, Hirokazu Ishii, Tetsuichi Wazawa, Joe Sakamoto, Yuichi Kozawa, Yuta Kato, Eri Mukai, Tomomi Nemoto, Takeharu Nagai

## Abstract

Two-photon excitation (2PE) microscopy reaches deep into tissue but remains diffraction-limited, and existing 2PE-STED and 2PE-RESOLFT schemes still rely on single-photon depletion or photoswitching. Here we present 2P-RESOLFT microscopy, in which excitation and reversible photoswitching are both two-photon-driven. Using the positive photoswitchable protein Padron2, we achieved 205-nm lateral resolution and resolved previously unreported membrane protrusions of β cells inside pancreatic pseudoislets.

## Main

Two-photon excitation (2PE) microscopy enables tissue imaging with intrinsic optical sectioning: nonlinear excitation confines the excitation volume to the focus and thereby suppresses the out-of-focus background that limits single-photon excitation microscopy^1,2^. However, its spatial resolution remains diffraction-limited. To overcome this limitation, 2PE microscopy has been combined with super-resolution techniques such as stimulated emission depletion (STED)^3,4,5^ and reversible saturable optical linear fluorescence transitions (RESOLFT)^6,7,8^, which use a donut-shaped beam to suppress fluorescence surrounding the excitation focus and confine emission to a sub-diffraction region. Nonetheless, 2PE-STED and 2PE-RESOLFT still rely on single-photon processes for stimulated emission depletion^4,5^ or photoswitching^8^, and thus forfeit the very features that make 2PE attractive: pinhole-free optical sectioning, deep penetration by near-infrared light, and reduced photodamage.

Here we conceived a fully two-photon-driven RESOLFT (2P-RESOLFT) scheme in which both the excitation and the photoswitching of a reversibly switchable fluorescent protein (rsFP)^7,9,10^ are driven by two-photon absorption. To test this concept, we first simulated 2P-RESOLFT for a generic positive-type green rsFP (p-rsGFP; see Methods for simulation details and Supplementary Note 1 for rsFP classification and selection). The 2P-RESOLFT illumination scheme with a p-rsFP therefore needs only two co-aligned beams, which illuminate the sample simultaneously and are scanned together as a single pattern, in contrast to the three-step illumination required for RESOLFT with negative-type rsFPs^11^. The beams are a 920 nm Gaussian beam that drives both two-photon ON-switching and excitation (2P ON-s/Ex) and a donut-shaped 780 nm beam for two-photon OFF-switching (2P OFF-s) of p-rsGFP (Fig. 1A, B). The simulated point spread functions (PSFs) show that the full width at half maximum (FWHM) is governed by both the intrinsic properties of the rsFP (Fig. 1C, D) and the imaging conditions (Fig. 1E, F). Briefly, the resolution improves as one increases the ratio of the intrinsic 2P OFF-s to ON-s coefficients of the rsFP (*k*_off,rsFP_/*k*_on,rsFP_) (Fig. 1C), fluorescence ON/OFF contrast (*F*_ON,920_/*F*_OFF,920_) (Fig. 1D), and the ratio of the 780 nm to 920 nm beam intensities (*A*_780,donut_/*A*_920,Gauss_) (Fig. 1E). In addition, at a constant *A*_780,donut_/*A*_920,Gauss_ ratio, higher beam intensities reduce the required pixel dwell time (*τ*_dwell_), whereas longer *τ*_dwell_ allows for the use of lower beam intensities while maintaining the spatial resolution (Fig. 1F), which affords considerably more flexibility in the choice of imaging conditions than STED microscopy. As with the beam intensities *A, k*_rsFP_ and *τ*_dwell_ exhibit a compensatory relationship (Extended Data Fig. 2). A detailed description of these simulation results is provided in Supplementary Note 2.

**Fig. 1.**
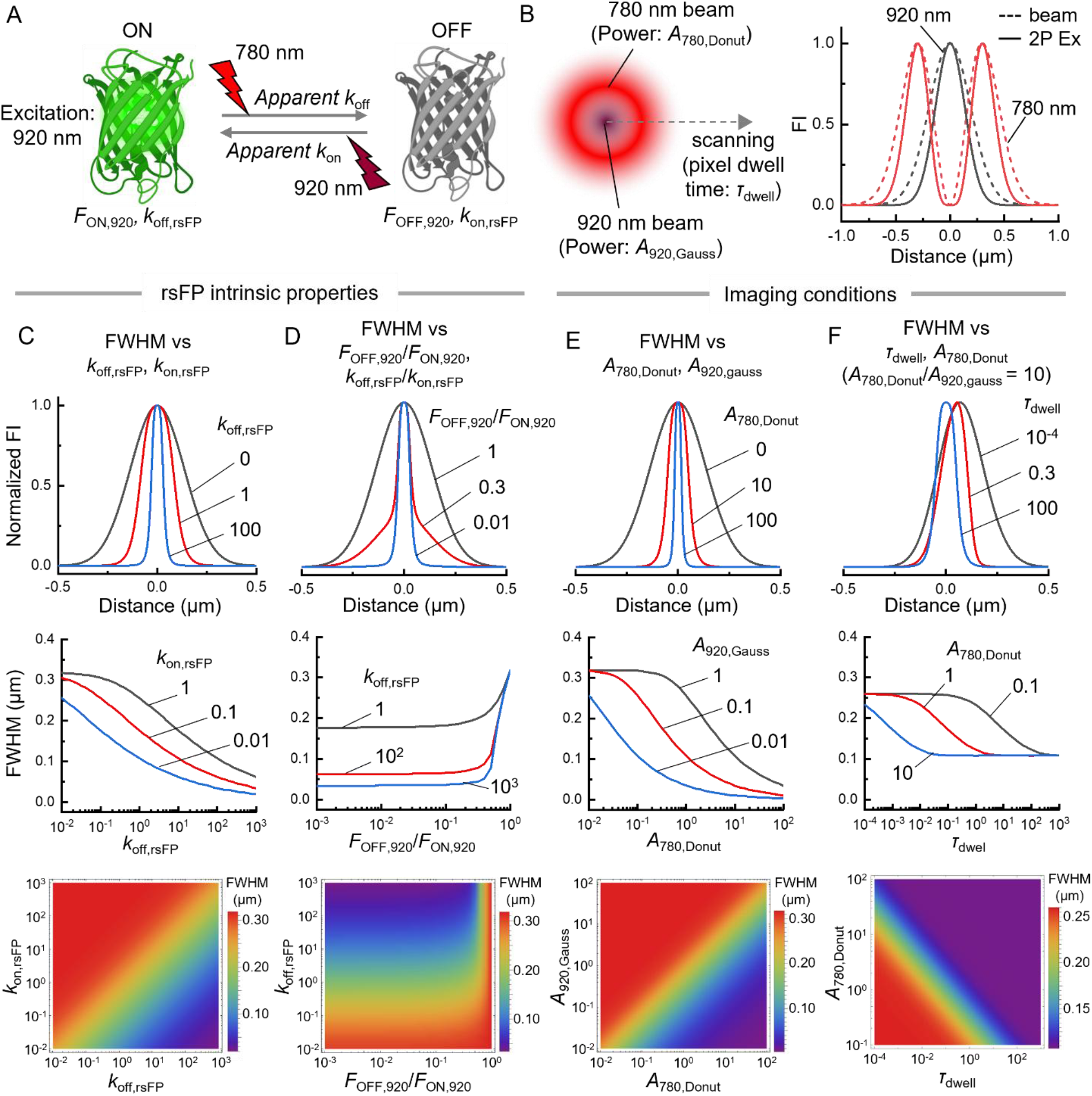
2P-RESOLFT simulation. (A) Photoswitching of a p-rsGFP. 780 nm for 2P OFF-s and 920 nm for 2P ON-s/Ex. *F*_ON,920_ and *F*_OFF,920_ are the fluorescence intensities of the ON and OFF states, respectively, excited by 920 nm. *k*_on,rsFP_ and *k*_off,rsFP_ are the rsFP intrinsic photoswitching coefficients, whereas the *k*_on,ap_ and *k*_off,ap_ are the photoswitching rate constants. (B) Illumination scheme of the 2P- RESOLFT microscope, with the beam and 2PE intensity profiles of the 920 nm Gaussian beam (black) and the 780 nm donut-shaped beam (red). (C–F) Simulated PSFs obtained with varying *k*_off,rsFP_, *k*_on,rsFP_ (C), *F*_ON,920_/*F*_OFF,920_ and *k*_off,rsFP_ (D), *A*_780,donut_, *A*_920,Gauss_ (E), and *A*_780,donut_ (at a constant intensity ratio *A*_780,donut_/*A*_920,Gauss_) and *τ*_dwell_ (F), the corresponding FWHM plotted against these parameters, and density maps of the FWHM. The simulation conditions are summarized in Supplementary Table 1. FI denotes fluorescence intensity.

To realize 2P-RESOLFT experimentally, we first characterized the photoswitching properties of the p-rsGFP Padron2^11^ in mammalian cells under 920 or 780 nm laser illumination (Fig. 2A). Both apparent *k*_on_ and *k*_off_ (*k*_on,ap_ and *k*_off,ap_, respectively) increased approximately quadratically with the intensity of the 920 nm and 780 nm beams, respectively, establishing that both ON- and OFF-switching of Padron2 are driven by two-photon processes (Fig. 2B). In addition, the *F*_ON,920_/*F*_OFF,920_ and *k*_off,rsFP_/*k*_on,rsFP_ ratios were determined to be 14.1 and 0.74, respectively (Fig. 2A and B).

**Fig. 2.**
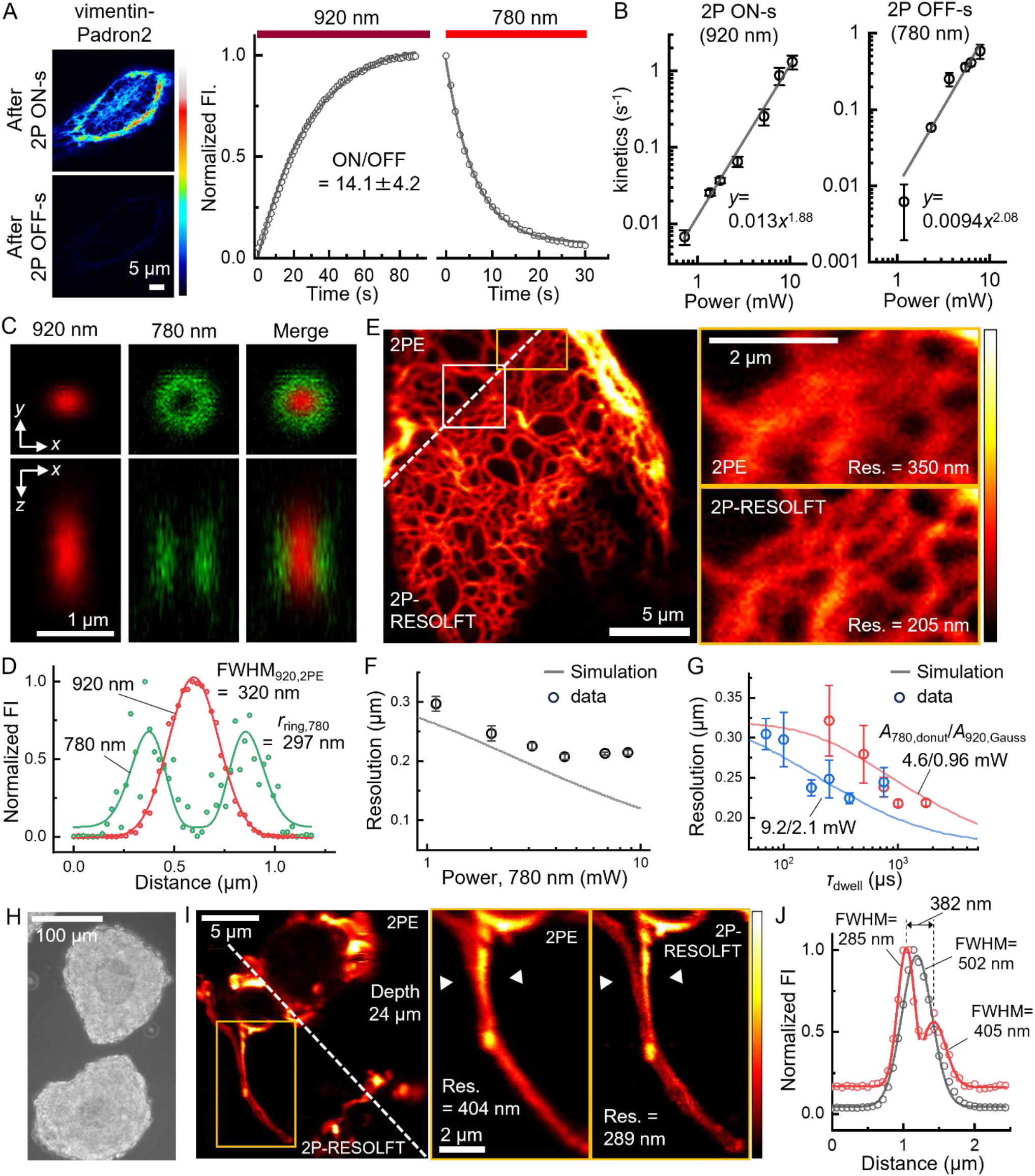
Implementation and demonstration of 2P-RESOLFT. (A) Representative 2PE image of a HeLa cell expressing vimentin-Padron2, and fluorescence intensity time courses under 920- and 780-nm illumination. ON/OFF contrast, mean ± SEM (*n*=4). (B) *k*_on,ap_ (left) and *k*_off,ap_ (right) plotted as a function of the 780 nm and 920 nm light intensity, respectively. Mean ± SEM (*n*=3-5 for each data point). (C, D) PSFs and the corresponding line profiles of 200 nm fluorescent beads under 920 nm Gaussian or 780 nm donut-shaped illumination. (E) 2PE and 2P-RESOLFT images of HeLa cell expressing vimentin-Padron2. Right: enlarged views of the orange-boxed region. White box: region shown in Extended Data Fig. 3A. (F) Simulated FWHM and spatial resolution plotted as a function of 780 nm laser intensities. Mean ± SEM (*n*=9 for each data point). (G) Simulated FWHM and spatial resolution plotted as a function of *τ*_dwell_ at the indicated 780/920 nm laser intensities (*A*_780,donut_/*A*_920,Gauss_). Mean ± SEM (*n*=9 for each data point). (H) Wide-field view of MIN6 pseudoislets. (I) 2PE and 2P- RESOLFT images of MIN6 cells expressing Padron2-CAAX within pseudoislets. (J) Normalized fluorescence intensity profiles along the line segments between the arrows in (I) (2PE, red; 2P- RESOLFT, black). Spatial resolutions (Res.) were determined by decorrelation analysis^12^. The observation conditions are summarized in Supplementary Table 2.

We then built a 2P-RESOLFT microscope. Notably, the two lasers need not be synchronized on the pulse timescale, because rsFP photoswitching proceeds from the ground state rather than from the short-lived excited state exploited in STED^3^. The FWHM of the 2PE PSF under the 920 nm Gaussian beam (FWHM_920,2PE_) and the ring radius of the donut-shaped 780 nm beam (*r*_ring,780_) were measured to be 320 nm and 297 nm, respectively (Fig. 2C, D). Image decorrelation analysis^12^ gave 350 nm for conventional 2PE microscopy and 205 nm for 2P-RESOLFT in HeLa cells expressing vimentin- Padron2 (Fig. 2E), a 1.7-fold gain that agrees with the representative line-profile FWHM of 206 nm and the simulated FWHM of 192 nm (Extended Data Fig. 3). Raising the 780 nm intensity sharpened the images further, although the gain saturated above 6.8 mW instead of continuing as predicted by the simulation (Fig. 2F), most likely because *k*_off,ap_ itself saturates under these conditions. As predicted by the simulation, the beam intensities primarily set the *τ*_dwell_ required to achieve the improved spatial resolution (Fig. 2G). Importantly, 2P-RESOLFT revealed fine, dim structures that conventional 2PE microscopy barely detected (Extended Data Fig. 4), reflecting efficient photon accumulation from weakly fluorescent structures at extended *τ*_dwell_, as in one-step RESOLFT^11^.

We next asked whether 2P-RESOLFT can image inside a biological tissue, and visualized the plasma membrane of pancreatic β cells within pseudoislets (Fig. 2H, I). 2P-RESOLFT delineated the plasma membrane of β cells within the pseudoislet more clearly than conventional 2PE microscopy (Extended Data Fig. 5A, B-1) and, in addition, resolved two thin membrane protrusions extending from a β cell that 2PE microscopy did not resolve (Fig. 2I, J, Extended Data Fig. 5A, B-2). To our knowledge, such thin membrane protrusions within pseudoislets have not been previously reported. 2P-RESOLFT thus resolves sub-diffraction architecture inside multicellular tissue and may open a window onto the structural basis of intercellular communication in pancreatic islets (see Supplementary Note 3 and Extended Data Fig. 5 for further details).

In summary, we have established fully two-photon-driven RESOLFT microscopy through numerical simulation, experimental validation, and imaging in a tissue model. Photon collection efficiency and axial resolution should improve further with non-descanned detection^2^ and with phase-engineering strategies that generate a three-dimensional OFF-switching distribution^13^, respectively. The performance of 2P-RESOLFT also depends critically on the rsFP itself. Further improvement of the photoswitching properties of existing p-rsFPs^11,14,15^ should sharpen the spatiotemporal resolution while lowering the required illumination intensity. Together, advances in instrumentation and rsFP engineering should broaden the reach of 2P-RESOLFT, opening the way to super-resolution imaging deep within living tissue.

## Methods

### 2P-RESOLFT simulation framework

In 2P-RESOLFT with p-rsFPs, the illumination scheme consists of a focused beam that induces both two-photon excitation and ON switching (2P ON-s/Ex), and a donut-shaped beam with zero intensity at the center that induces two-photon OFF switching (2P OFF-s) (Fig. 1A, B). Upon superimposing these beams and scanning them across the sample, p-rsFPs in the region exposed to the 2P OFF-s beam are driven into the OFF state and no longer emit fluorescence under 2P excitation. As a result, fluorescence emission is confined to the central region of the donut, where the 2P OFF-s beam intensity is minimal.

In the simulation of 2P-RESOLFT microscopy, we assumed a p-rsGFP as the fluorescent probe, a 780 nm pulsed laser for 2P OFF-s, and a 920 nm pulsed laser for 2P ON-s/Ex (Fig. 1A). In STED microscopy, stimulated emission must occur within the fluorescence lifetime of the excited state (typically a few nanoseconds), requiring sub-nanosecond temporal synchronization between the excitation and depletion pulses. In contrast, photoswitching in rsFPs is initiated from the ground state and is therefore not constrained by the nanosecond lifetime of the excited state. Therefore, precise temporal synchronization between the Gaussian and donut beams is not required for 2P-RESOLFT. Although pulsed lasers were used experimentally, both were treated as pseudo-continuous-wave (CW) sources in the simulations because the relevant switching dynamics are expected to depend primarily on accumulated photon dose rather than pulse timing. The normalized intensity profiles of the 920 nm Gaussian beam for 2P ON-s/Ex and of the donut-shaped 780 nm beam for 2P OFF-s were simulated using the following equations:

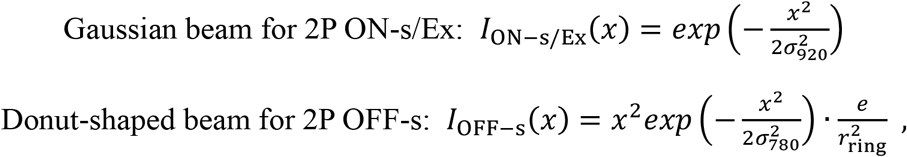

where *x* is the lateral distance from the beam center, *e* is the base of the natural logarithm, and *σ*_920_ is a constant related to the full width at half maximum of the intensity profile of the 920 nm Gaussian beam (FWHM_920,1PE_) by

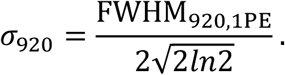

FWHM_920,1PE_ >was estimated to be 452 nm from the experimentally determined FWHM_920,2PE_ (Fig. 2C, D) using the relationship FWHM_920,1PE_ = √ 2 × FWHM_920,2PE_, which holds because the two-photon excitation probability scales with the square of the beam intensity. The parameter *σ*_780_ is defined as

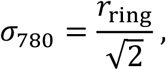

where *r*_ring_ >represents the experimentally measured radius at which the donut beam reaches its maximum intensity (*r*_ring_ = 297 nm) (Fig. 2C, D). The Donut-shaped beam profile is normalized by 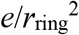 so that the profile peaks at unity at *x* = *r*_ring_.

A point-like rsFP was assumed to be located at the center, and the illumination beams were scanned from left to right at a constant velocity. The beam center position is expressed as

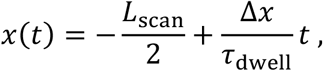

where *L*_scan_ >is the one-dimensional line scan length, *τ*_dwell_ is the pixel dwell time, and Δ*x* is the pixel size. The rsFP undergoes reversible photoswitching between the ON and OFF states upon illumination with the 920 and 780 nm beams (Fig. 1A). The population dynamics of the ON and OFF states are described by the following differential equations with the initial condition:

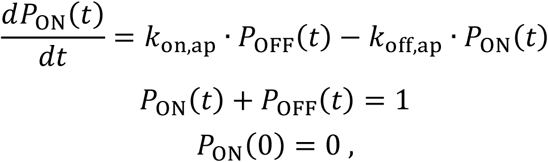

where *P*_ON_(*t*) and *P*_OFF_(*t*) are the mole fractions of the ON and OFF states, respectively, and apparent *k*_on_ and *k*_off_ (*k*_on,ap_ and *k*_off,ap_, respectively) are the rate constants of 2P ON- and 2P OFF-switching, respectively (Fig. 1A), defined as

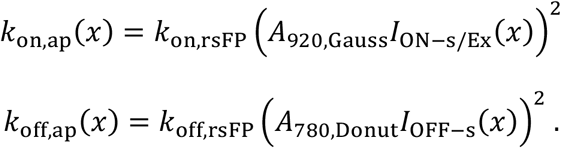

*k*_on,rsFP_ and *k*_off,rsFP_ >are the 2P ON- and OFF-switching rate coefficients at the beam peak, respectively (Fig. 1A). These coefficients incorporate the intrinsic photoswitching properties of the rsFP, including the photoswitching quantum yield and the two-photon absorption cross section. *A*_920,Gauss_ and *A*_780,donut_ are scaling factors that determine the peak excitation intensity of the 920 nm Gaussian beam and the 780 nm donut beam, respectively (Fig. 1B). Under 920 and 780 nm illumination, both the ON and OFF states are excitable and capable of generating fluorescence. However, fluorescence induced by the 780 nm beam was assumed to be negligible as

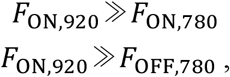

where *F*_*x,y*_ is fluorescence intensity from state *x* (ON or OFF) under excitation at wavelength *y* (920 or 780 nm). Therefore, the detected fluorescence was approximated by that generated under 920 nm two-photon excitation. The fluorescence signal emitted during the *τ*_dwell_ at each scan position *x*(*t*) is calculated as

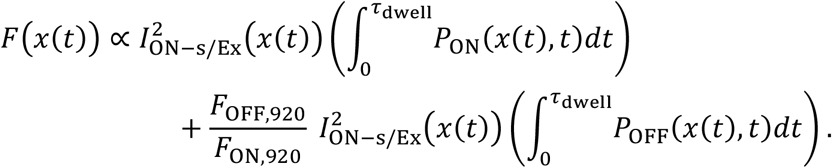

For analysis, the simulated fluorescence profile *F*(*x*) was normalized to its peak intensity. The FWHM of the PSF was defined as the distance between the outermost left and right positions at which the normalized intensity fell to 0.5. The FWHM of the resulting PSF was calculated as a function of *k*_on,rsFP_ and *k*_off,rsFP_ (Fig. 1C). To investigate the effect of the two-photon ON/OFF contrast, the FWHM was calculated as a function of the *k*_off,rsFP_ and *F*_OFF,920_/*F*_ON,920_ (Fig. 1D). The dependence of the FWHM on the excitation intensities *A*_920,Gauss_ and *A*_780,donut_ was examined in Fig. 1E. In Fig. 1F, the FWHM was calculated as a function of *τ*_dwell_ and the excitation intensities *A*_780,donut_, while maintaining a constant intensity ratio (*A*_780,donut_/*A*_920,gauss_ = 10).

In Fig. 2F and Extended Data Fig. 3C, simulated FWHMs were calculated based on the experimental conditions. The definitions of the intrinsic rsFP photoswitching coefficients, *k*_rsFP_, and the illumination parameter, *A*, used in the simulations differ from those of the corresponding experimentally determined values, resulting in a discrepancy between the experimental and simulated values. However, at sufficiently long *τ*_dwell_, the spatial resolution is primarily determined by the ratios *k*_off,rsFP_/*k*_on,rsFP_ and *A*_780,donut_/*A*_920,Gauss_ (Fig. 1C, E, and Supplementary Note 2). We therefore calculated the FWHM using these experimentally obtained ratios. The *k*_off,rsFP_/*k*_on,rsFP_ ratio was determined from Fig. 2B, while the illumination parameters were determined from the laser powers measured just above the objective aperture using a power meter (PM100D, Thorlabs) with a sensor (S170C, Thorlabs). In Fig. 2G, the spatial resolution as a function of *τ*_dwell_ depends not only on the *k*_off,rsFP_>/*k*_on,rsFP_> and *A*_780,donut_/*A*_920,Gauss_ but also on the individual values of *k*_rsFP_ and *A* (Fig. 1F and Extended Data Fig. 2). We therefore estimated *k*_rsFP_ by least-squares fitting of the simulated FWHM to the experimentally measured resolutions while maintaining a constant *k*_off,rsFP_/*k*_on,rsFP_ ratio of 0.74. The simulation conditions are summarized in Supplementary Table 1. These simulations were performed using Wolfram Mathematica 15.0.1.0 software (Wolfram Research).

### Gene construction

For the vimentin-Padron2 construct, the Padron2 gene (derived from Addgene #172355) was amplified by PCR (KOD plus NEO; TOYOBO) using primers containing BamHI and EcoRI restriction sites and used to replace the rsZACRO gene in vimentin-rsZACRO pcDNA3^15^. For the Padron2-CAAX construct, the Padron2 gene was amplified by PCR using primers containing HindIII and EcoRI restriction sites and used to replace the rsZACRO gene in rsZACRO-CAAX pcDNA3.

### Mammalian cell culture and transfection

HeLa cells, provided by the RIKEN Cell Bank through the National BioResource Project (RCB0007), were cultured in Dulbecco’s modified Eagle’s medium (D-MEM (low glucose), 041-29775, Wako) supplemented with 10% (v/v) fetal bovine serum in a CO_2_ incubator at 37 °C with 5% CO_2_. HeLa cells transferred to a 35 mm glass-bottom dish were transfected using polyethylenimine PEI Max 40K (Polysciences) or Lipofectamine LTX and Plus (Invitrogen) according to the manufacturer’s instructions. More than 24 h after transfection, the medium was replaced with DMEM/F12 (11039021; Thermo Fisher Scientific) supplemented with HEPES, or DMEM fluoroBrite (Gibco) supplemented with HEPES (Gibco) and GlutaMax (Gibco) before imaging.

MIN6 cells, a cell line derived from mouse pancreatic β cells, were cultured in D-MEM (high-glucose) (043-30085, Wako) supplemented with 10% fetal bovine serum (CORNING), 50 U/mL penicillin– streptomycin (Gibco), and 35 µM β-mercaptoethanol (Gibco) in a CO_2_ incubator at 37 °C with 5% CO_2_. The culture medium was replaced every 2 days.

To prepare pseudoislets, MIN6 cells were suspended to a density of 2.5 × 10^4^ cells/mL in culture medium. Aliquots (20 µL) of the cell suspension were placed as droplets on the lids of 10 cm culture dishes and incubated for 3 days in a CO_2_ incubator using the hanging-drop method. Subsequently, the formed pseudoislets were detached from the lids with culture medium and transferred to 10 cm dishes coated with collagen (Cell Matrix Type I-P, Nitta Gelatin Inc.). The pseudoislets were further cultured for 2 to 4 days. MIN6 cells were transfected with Lipofectamine LTX and Plus according to the manufacturer’s instructions, starting during hanging-drop preparation. The transfection complexes were also added to the culture medium throughout the subsequent culture period until immediately before imaging. The medium was replaced with DMEM fluoroBrite supplemented with HEPES and GlutaMax before imaging.

### Optical setup for 2P-RESOLFT microscope

The 2P-RESOLFT microscope was built on an inverted microscope (ECLIPSE Ti2-U, Nikon) equipped with two femtosecond pulsed light sources and a spatial light modulator (SLM; SLM-200, Santec). The two-photon excitation sources for fluorescence and photoswitching were a mode-locked Ti:sapphire femtosecond laser operating at 780 nm (Tsunami, Spectra-Physics) and a mode-locked femtosecond fiber laser operating at 920 nm (Alcor 920-1, Spark Lasers). Unlike STED microscopy, precise temporal synchronization of the excitation and OFF-switching laser pulses is not required because photoswitching of rsFPs is initiated from the ground state and is therefore not constrained by the nanosecond lifetime of the excited state. Therefore, the two pulsed lasers were operated at a repetition rate of 80 MHz without synchronization between them.

To generate the donut-shaped 780 nm beam, the SLM was introduced into a dedicated optical path for the 780 nm laser. The objective back pupil was relayed onto the SLM using a 4f optical system, and a vortex phase pattern was applied to the 780 nm beam. Phase patterns for correcting astigmatism, coma, and trefoil aberrations were superimposed on the vortex phase pattern to obtain a rotationally symmetric donut-shaped intensity distribution. A custom-developed transmissive liquid-crystal device (tLCD-P)^16,17,18^ was inserted between the objective turret and the objective lens to convert the vortex beam to circular polarization, thereby preserving the central intensity minimum of the donut-shaped beam.

The combined 780 nm and 920 nm beams were raster-scanned across the sample using a two-axis galvo-mirror scanner (8315KM40B, Cambridge Technology). The beams were focused through a 60× silicone-immersion objective lens (CFI Plan Apo Lambda S 60XC Sil, NA 1.30, Nikon). Fluorescence was descanned and separated from the excitation light using a dichroic mirror (Di03-R561-t1, Semrock) and an appropriate emission filter (XF3079 535AF26, Omega Optical). The signal was then directed through a confocal pinhole module (MPH16, Thorlabs) and detected with a GaAsP photomultiplier tube (PMT; H10771PA-40, Hamamatsu Photonics). The pinhole size was set to 300 µm. This setting provided modest rejection of out-of-focus background signal while limiting fluorescence loss^19^.

### Measurement of 2P photoswitching ON/OFF contrast and kinetics

The 2P photoswitching ON/OFF contrasts of rsEGFP2, rsGamillus-S, and Padron2 were characterized by observing HeLa cells expressing LifeAct-rsEGFP2, LifeAct-rsGamillus-S, and vimentin-Padron2 using a two-photon excitation microscope (A1R-MP+, Nikon), with a water-immersion objective lens (Nikon, Apo LWD 25×/1.10 NA). A Ti:sapphire laser (MaiTai eHP DeepSee, Spectra-Physics; repetition rate: 80 MHz) was used as the excitation laser light source at wavelengths of 920 nm and 780 nm. The laser power measured immediately below the objective lens was 3.9 mW at 920 nm and 3.5 mW at 780 nm. Fluorescence signals were detected using non-descanned detectors equipped with GaAsP photomultiplier tubes, with no optical filters placed in front of the detectors. The changes in fluorescence intensity were fitted to the following function,

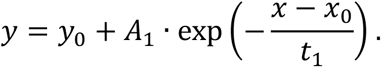

where *y*_0_ >is the baseline fluorescence intensity, *A*_1_ represents the initial deviation from the baseline fluorescence intensity, *x*_0_ is the initial time point, and *t*_1_ is the time constant. For n-rsFPs, the ON/OFF contrast was calculated as the ratio of the initial fluorescence intensity (*y*_0_+*A*_1_) to the asymptotic fluorescence intensity (*y*_0_). For Padron2, the ON/OFF contrast was calculated as the ratio of the asymptotic fluorescence intensity (*y*_0_) to the initial fluorescence intensity (*y*_0_+*A*_1_).

Power-dependent 2P ON- and OFF-switching kinetics of Padron2 (Fig. 2B) were characterized by observing HeLa cells expressing vimentin-Padron2 using the 2P-RESOLFT microscope described above, with 780 nm and 920 nm Gaussian beams. To determine the photoswitching rate constants of Padron2, fluorescence intensity time courses were analyzed using pseudo-first-order kinetics, following previously described methods^20^. These fitting analyses were performed using OriginPro 2021 (OriginLab).

### Image analysis for spatial resolutions and length of observed structures in mammalian cells

The imaging data were analyzed using ImageJ Fiji software^21^. The spatial resolutions were analyzed by image decorrelation analysis using an ImageJ plugin (available at https://github.com/Ades91/ImDecorr.git), which estimates the spatial resolution of individual images based on the correlation between the input image and its normalized Fourier transform and shows good agreement with FWHM-based measurements^12^. The FWHMs were determined by fitting Gaussian functions to the intensity line profiles. The two-photon donut PSF was fitted with 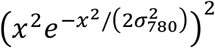. The fitted parameter σ_780_> was then used to define the donut beam profile, where the ring radius is given by 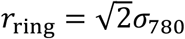 (see “2P-RESOLFT simulation framework” section in Methods). These fitting analyses were performed using OriginPro 2021 (OriginLab). The lengths of the observed structures in MIN6 pseudoislets were measured with the Neuroanatomy SNT, an ImageJ plugin.

## Data availability

The authors declare that the data supporting the findings of this study are available within the paper and its Supplementary Information files. Should any raw data files be needed in another format they are available from the corresponding author upon reasonable request.

## Code availability

The Wolfram Mathematica code for the PSF and FWHM calculations is available from the corresponding author upon reasonable request during peer review and will be made publicly available upon publication.

## Acknowledgements

This work was in part supported by grants from Core Research for Evolutionary Science and Technology, Japan Science and Technology Agency (JPMJCR15N3 and JPMJTR254C to T. Nagai); JST SPRING (JPMJSP2101 to Y. Kato); Japan Society for the Promotion of Science (18H05410, 21K19225, 23H05421, 25H02457, 26K21746 to T. Nagai; 22H04926, Advanced Bioimaging Support (ABiS), 25H01025 to T. Nemoto; 22K04891 to T. Wazawa; 26K17749 to J. Sakamoto; 25K10607 to H. Ishii; 26K17746, 26K03263 to R. Ozaki-Noma); AMED Brain/MINDS (19dm0207078) and Brain/MINDS 2.0 (24wm0625105) to T. Nemoto; the Cooperative Study Program (24NIPS243, 25NIPS255, 26NIPS255 to T. Nagai) of National Institute for Physiological Sciences; Joint Research of Exploratory Research Center on Life and Living Systems (ExCELLS) (23EXC601 and 25EXC602 to T. Nemoto); ExCELLS Encouragement Research for Young Scientists (26-Y5 to R. Ozaki-Noma); Toyoaki Scholarship Foundation to H. Ishii.

## Author information

Authors and Affiliations

**SANKEN, The University of Osaka, 8-1 Mihogaoka, Ibaraki, Osaka 567-0047, Japan**

Ryohei Ozaki-Noma, Tetsuichi Wazawa, Takeharu Nagai

**School of Life Science, The Graduate University for Advanced Studies, SOKENDAI, Okazaki, Aichi 444-8787, Japan**

Hirokazu Ishii, Joe Sakamoto, Tomomi Nemoto

**Division of Biophotonics, National Institute for Physiological Sciences (NIPS), National Institutes of Natural Sciences (NINS), Okazaki, Aichi 444-8585, Japan**

Hirokazu Ishii, Joe Sakamoto, Tomomi Nemoto

**Biophotonics Research Group, Exploratory Research Center on Life and Living Systems (ExCELLS), NINS, Okazaki, Aichi 444-8787, Japan**

Hirokazu Ishii, Joe Sakamoto, Tomomi Nemoto

**Transdimensional Life Imaging Division, Institute for Open and Transdisciplinary Research Initiatives, The University of Osaka, Suita, Osaka 565-0871, Japan**

Tetsuichi Wazawa, Takeharu Nagai

**Department of Pharmacology, Hamamatsu University School of Medicine, Hamamatsu, Shizuoka 431-3192, Japan**

Joe Sakamoto

**Institute of Multidisciplinary Research for Advanced Materials, Tohoku University, Sendai, Miyagi 980-8577, Japan**

Yuichi Kozawa

**Department of Biomedical Sciences, College of Life Sciences, Ritsumeikan University, 1-1-1 Nojihigashi, Kusatsu, Shiga 525-8577, Japan**

Yuta Kato, Eri Mukai

## Contributions

Ryohei Ozaki-Noma: Conceptualization (lead), Methodology (supporting), Investigation (lead), Software (supporting), Resources (equal), Writing–original draft (lead), Writing–Review & Editing (equal), Visualization (lead). Hirokazu Ishii: Conceptualization (supporting), Methodology (lead), Investigation (supporting), Writing–Original Draft (supporting), Writing–Review & Editing (equal). Tetsuichi Wazawa: Investigation (supporting), Software (lead), Writing–Review & Editing (equal). Joe Sakamoto: Conceptualization (supporting), Methodology (supporting), Resources (supporting), Writing–Review & Editing (equal). Yuichi Kozawa: Methodology (supporting), Writing–Review & Editing (equal). Yuta Kato: Resources (equal), Writing–Review & Editing (equal). Eri Mukai: Resources (supporting), Writing–Review & Editing (equal). Tomomi Nemoto: Conceptualization (supporting), Writing–Review & Editing (equal), Supervision (equal). Takeharu Nagai: Conceptualization (supporting), Writing–Review & Editing (lead), Supervision (equal).

---

**Extended Data Fig. 1.**
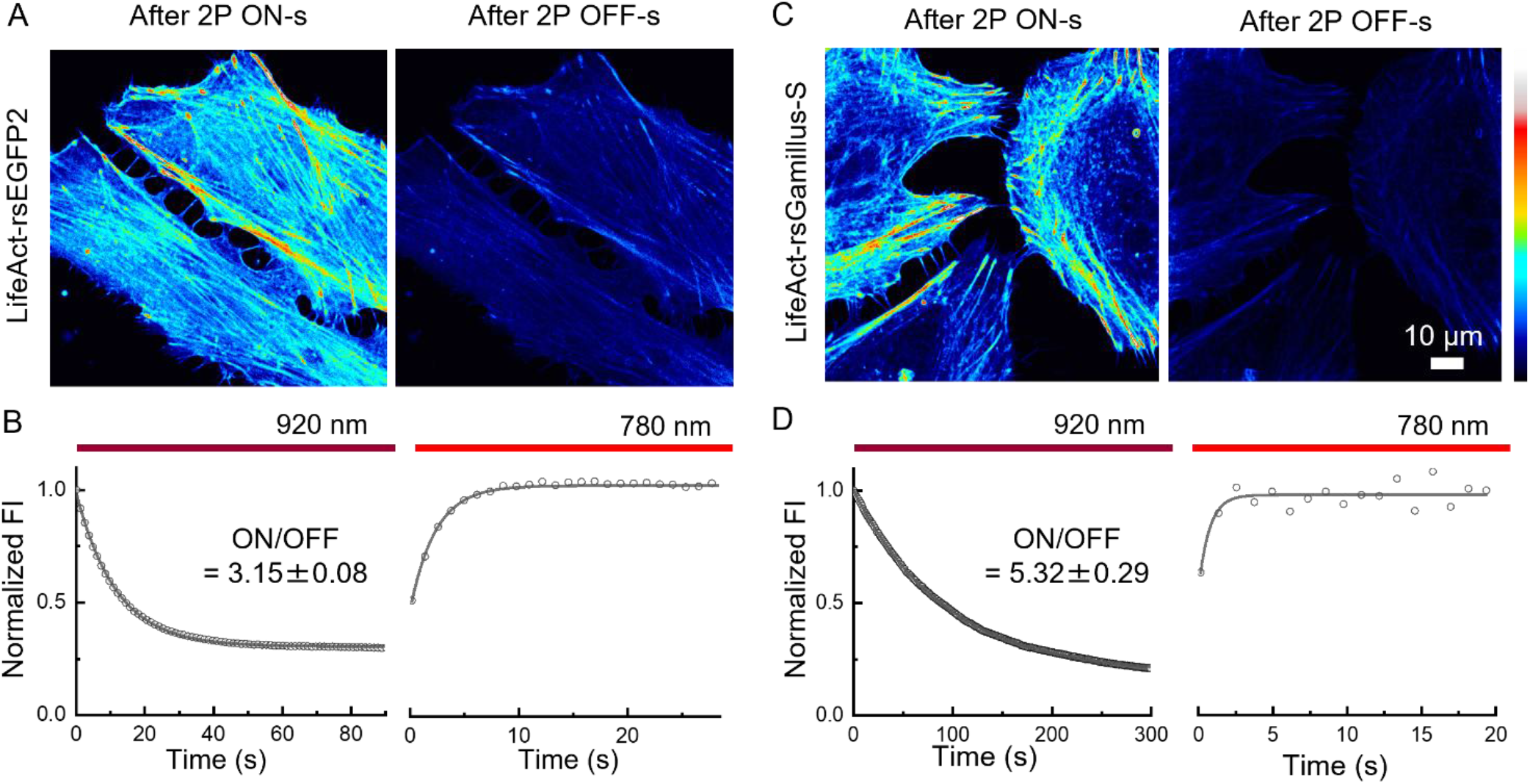
Two-photon photoswitching of n-rsFPs. (A, C) Representative fluorescence images of HeLa cells expressing LifeAct-rsEGFP2 and LifeAct- rsGamillus-S, respectively. (B, D) Representative time courses of the fluorescence intensity of rsEGFP2 and rsGamillus-S, respectively, in HeLa cells during photoswitching under 920 nm for 2P OFF-s and 780 nm illumination for 2P ON-s. These ON/OFF contrasts were determined as the ratio of the asymptotic fluorescence intensity obtained by fitting (see **Methods** section) the OFF-switching kinetics to the initial fluorescence intensity. Mean ± SEM (*n*=6 for rsEGFP2 and *n*=4 for rsGamillus- S). FI denotes fluorescence intensity.

**Extended Data Fig. 2.**
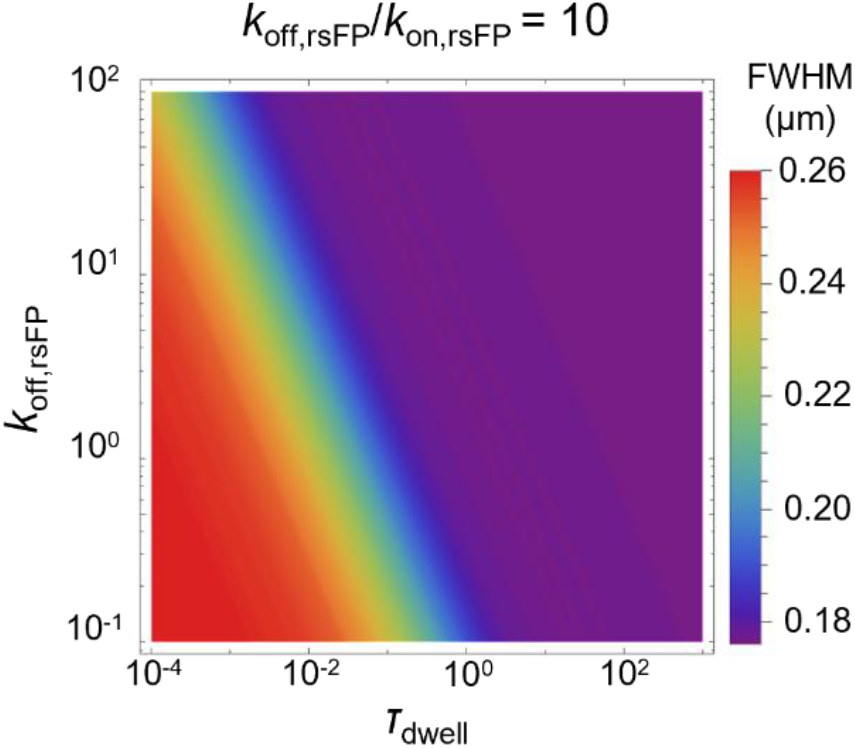
A density map of the FWHM as a function of *k*_off,rsFP_ (at a constant ratio *k*_off,rsFP_/*k*_on,rsFP_) and *τ*_dwell_. Constant-resolution contours along diagonal lines with negative slopes indicate that the rsFP intrinsic photoswitching coefficients and *τ*_dwell_ compensate for each other.

**Extended Data Fig. 3.**
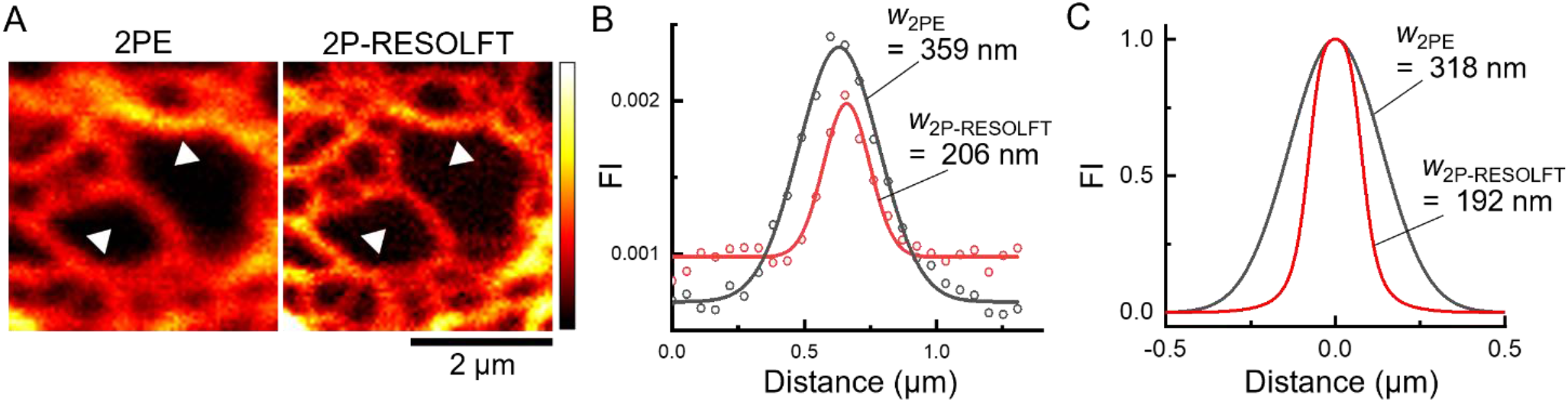
FWHM of 2P-RESOLFT imaging and simulation. (A) Enlarged view of the region enclosed by the white box in Fig. 2E by 2PE and 2P-RESOLFT microscope. (B) Representative line profiles of fluorescence intensity for 2P (black) and 2P-RESOLFT (red) along the line segments indicated by arrows in A. Gaussian distribution functions fitted to the data points. (C) Simulated PSF for 2P (black line) and 2P-RESOLFT (red line) using experimentally determined parameters of *k*_off,rsFP_/*k*_on,rsFP_, *F*_ON,920_/*F*_OFF,920,_ and *A*_780,donut_/*A*_920,Gauss_ (Supplementary Table 1). *w* indicate FWHM (nm) of the peaks.

**Extended Data Fig. 4.**
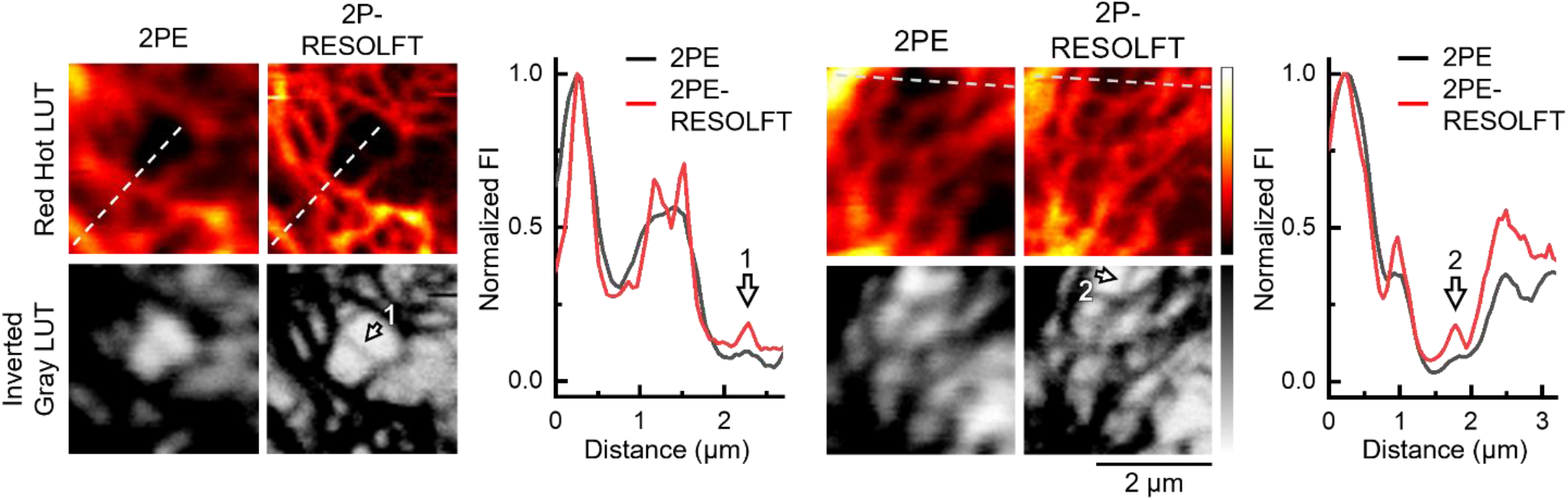
Visualization of dim, fine structures by 2PE and 2P-RESOLFT imaging. Representative images of HeLa cells expressing vimentin-Padron2 acquired by 2PE and 2P-RESOLFT microscopy, displayed using Red Hot and inverted Gray LUTs, respectively, with intensity line profiles along the dashed lines indicated in the images. The arrows indicate structures that are poorly visualized by conventional 2PE microscopy under identical imaging conditions.

**Extended Data Fig. 5.**
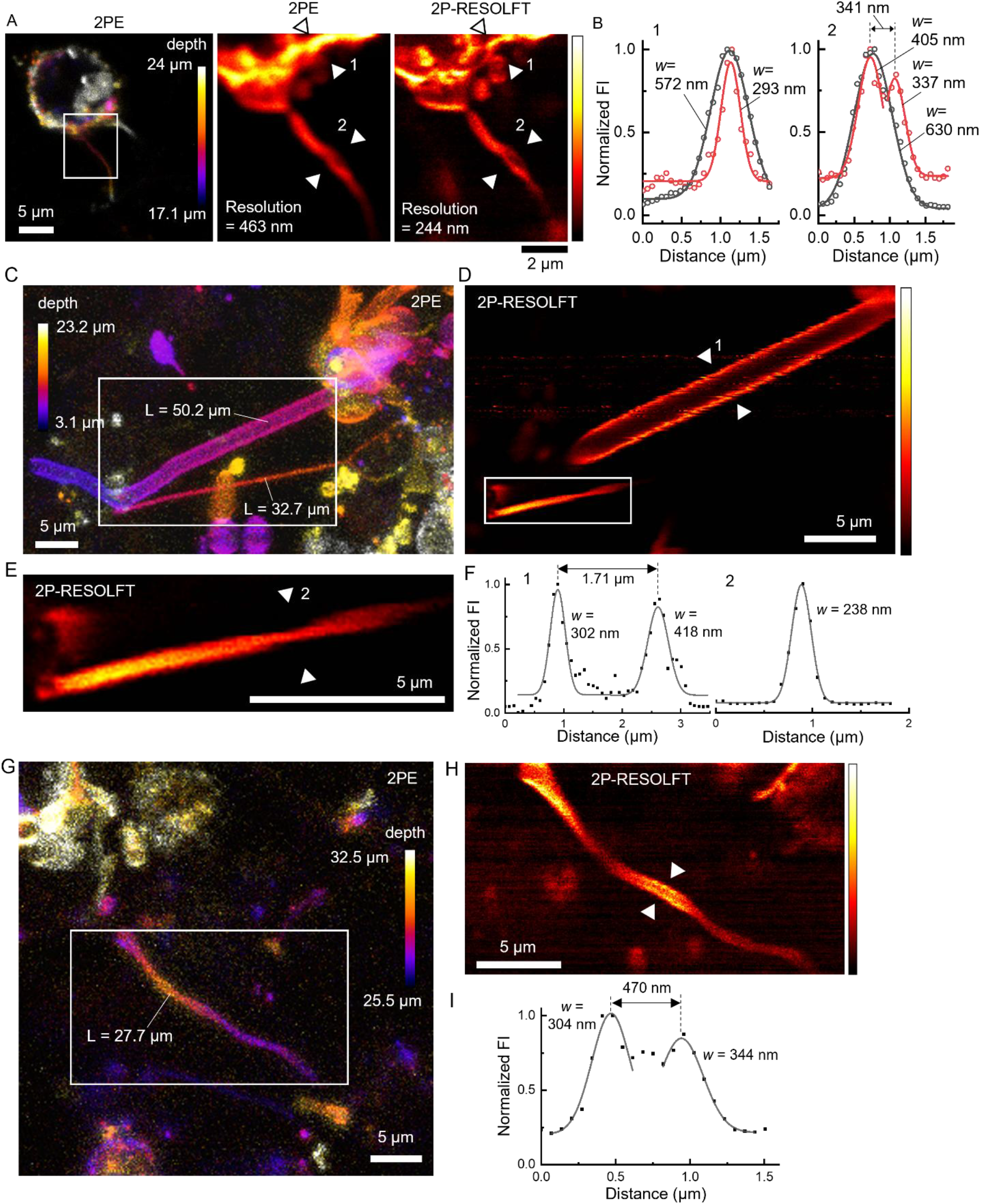
Membrane protrusions of MIN6 cells in pseudoislets. (A) Left: Fluorescence images of MIN6 cells expressing Padron2-CAAX in pseudoislets, acquired with 2PE microscopy. Middle and right: The images show an enlarged view of the region enclosed by the white box, acquired with 2PE and 2P-RESOLFT microscopy. Spatial resolutions determined by decorrelation analysis are indicated. (B) Line profiles of fluorescence intensity for 2PE (black) and 2P-RESOLFT (red) along the line segments between the arrows in (A). (C, G) Fluorescence images of MIN6 cells expressing Padron2-CAAX in pseudoislets, acquired with 2PE microscopy. (D, H) Magnified 2P-RESOLFT views of the area enclosed by squares in (C) and (G), respectively. (E) A magnified 2P-RESOLFT view of the area enclosed by squares in (D). (F) Line profiles of fluorescence intensity along the line segments between the arrows in (D) and (E). (I) Line profiles of normalized fluorescence intensity along the line segments between the arrows in (H). *w* values indicate FWHM (nm).

