## Supplementary Notes for "Fully Two-Photon-Driven RESOLFT Microscopy for Super-Resolution Imaging inside Tissue"

**Supplementary Note 1. Reversibly photoswitchable fluorescent proteins for 2P-RESOLFT**

Reversibly switchable fluorescent proteins (rsFPs) are genetically encodable fluorescent proteins that can be reversibly switched between a fluorescent ON state and a dim OFF state by illumination with specific wavelengths. rsFPs are classified as negative- or positive-type according to the direction of the photoswitching that is triggered by the fluorescence excitation light. In negative-type rsFPs (n-rsFPs), the excitation wavelength also induces OFF switching, whereas ON switching is triggered by a different wavelength[5,9,10]. For example, Dronpa[22] undergoes fluorescence excitation and OFF switching upon 488 nm illumination and is switched back to the ON state by 405 nm illumination. In contrast, positive-type rsFPs (p-rsFPs) undergo fluorescence excitation and ON switching at the same wavelength, while OFF switching is induced by a different wavelength. For example, Padron[23] is excited and switched ON by 488 nm illumination and switched OFF by 405 nm illumination.

For the implementation of fully two-photon-driven RESOLFT (2P-RESOLFT) microscopy, two critical issues related to the photoswitching properties of rsFPs must be addressed: (i) whether reversible photoswitching can be efficiently induced by two-photon excitation and (ii) whether the resulting ON/OFF contrast is sufficient for improvement of spatial resolution in 2P-RESOLFT. The ON/OFF contrast, in particular, is an intrinsic property of the rsFP that cannot readily be improved by optimizing the imaging conditions, and it is therefore a key determinant of 2P-RESOLFT performance. Furthermore, the illumination scheme required for RESOLFT microscopy depends on whether the rsFP is of the negative- or the positive-type. For n-rsFPs, 2P-RESOLFT requires a three-step illumination sequence: (1) Gaussian-shaped excitation for ON-switching, (2) donut-shaped excitation for OFF-switching, and (3) Gaussian-shaped excitation to detect the fluorescence from the remaining ON-state n-rsFP molecules. In contrast, for p-rsFPs, the RESOLFT illumination scheme can be simplified to a single step by spatially overlapping and scanning the two beams, a Gaussian beam for ON-switching/excitation and a donut-shaped beam for OFF-switching[11].

To address these questions, we evaluated the two-photon photoswitching properties of n-rsFPs, rsEGFP2, a well-established rsFP for RESOLFT microscopy[24] (Extended Data Fig. 1A, B), rsGamillus-S[25], an rsFP previously applied to super-resolution imaging based on two-photon ON-switching[26] (Extended Data Fig. 1C, D), and the p-rsFP Padron2[11] (Fig. 2A). Under single-photon excitation, rsEGFP2, rsGamillus-S, and Padron2 exhibited an ON/OFF contrast of 52, 103[25], and 53[11], respectively. However, under two-photon excitation using 780 nm and 920 nm, rsEGFP2 and rsGamillus-S showed ON/OFF contrasts of only 3.15 and 5.32, respectively (Extended Data Fig. 1B, D). This discrepancy between the single-photon and two-photon contrasts is most likely attributable to spectral crosstalk between the ON and OFF states under 780 and 920 nm two-photon excitation, resulting in incomplete photoswitching. Whereas the two-photon ON/OFF contrasts of the n-rsFPs tested were insufficient, Padron2 provided a contrast of 14.1, high enough for a substantial gain in resolution in 2P-RESOLFT microscopy (Fig. 1D and 2A). Padron2 was engineered to minimize residual fluorescence in the OFF state and to improve resistance to photoswitching fatigue, and it has been used successfully in one-step RESOLFT microscopy[11]. We therefore selected Padron2 and adopted the single-step RESOLFT illumination scheme for the present 2P-RESOLFT implementation.

**Supplementary Note 2. PSFs and FWHMs of 2P-RESOLFT simulation**

Details of the calculation procedure and simulation conditions in the 2P-RESOLFT simulation are described in "2P-RESOLFT simulation framework" section in Methods and Supplementary Table 1. We first examined the dependence of the simulated FWHM on the intrinsic properties of the rsFP, including the photoswitching rate coefficients ($k_{\mathrm{off,rsFP}}$ and $k_{\mathrm{on,rsFP}}$) and ON/OFF contrast ($F_{\mathrm{ON,920}}/F_{\mathrm{OFF,920}}$).

In the upper panel of Fig. 1C, simulated PSFs are shown for a fixed $k_{\mathrm{on,rsFP}} = 0.1$ and varying $k_{\mathrm{off,rsFP}}$ values of 0, 1, and 100. The case of $k_{\mathrm{off,rsFP}} = 0$ corresponds to conventional two-photon excitation, yielding a PSF with an FWHM of 318 nm. As $k_{\mathrm{off,rsFP}}$ increases, the PSF becomes narrower, resulting in a corresponding decrease in the FWHM, as shown in the middle panel of Fig. 1C. In addition, a density map of the FWHM as a function of $k_{\mathrm{off,rsFP}}$ and $k_{\mathrm{on,rsFP}}$ reveals that the FWHM is primarily determined by the ratio $k_{\mathrm{off,rsFP}}/k_{\mathrm{on,rsFP}}$, rather than the individual rate constants, as indicated by the constant-FWHM contours aligned along diagonal lines. In these simulations, the pixel dwell time ($\tau_{\mathrm{dwell}}$) was set sufficiently long, with $F_{\mathrm{OFF,920}}/F_{\mathrm{ON,920}}$ set to 0 and $A_{\mathrm{780,donut}}$ and $A_{\mathrm{920,Gauss}}$ set to 1 (Supplementary Table 1).

In Fig. 1D, the upper panel shows simulated PSFs for $F_{\mathrm{OFF,920}}/F_{\mathrm{ON,920}}$ values of 1, 0.3, and 0.01. The case of $F_{\mathrm{OFF,920}}/F_{\mathrm{ON,920}} = 1$ corresponds to a complete absence of ON/OFF contrast, that is, to conventional two-photon excitation. Compared with $F_{\mathrm{OFF,920}}/F_{\mathrm{ON,920}} = 0.01$, the PSF for $F_{\mathrm{OFF,920}}/F_{\mathrm{ON,920}} = 0.3$ exhibits broader side lobes, which degrade the spatial resolution. Consistently, the FWHM decreased with decreasing $F_{\mathrm{OFF,920}}/F_{\mathrm{ON,920}}$, and the middle panel and the density map show that the FWHM reaches a plateau once $F_{\mathrm{OFF,920}}/F_{\mathrm{ON,920}}$ falls below approximately 0.1, indicating that an ON/OFF contrast greater than 10 is desirable for a p-rsGFP under two-photon photoswitching. In these simulations, $\tau_{\mathrm{dwell}}$ was set sufficiently long, with $A_{\mathrm{780,donut}}$ and $A_{\mathrm{920,Gauss}}$ set to 1 (Supplementary Table 1).

Next, we examined the dependence of the simulated FWHM on the imaging parameters, including beam intensities ($A_{\mathrm{780,donut}}$ and $A_{\mathrm{920,Gauss}}$) and $\tau_{\mathrm{dwell}}$. In the upper panel of Fig. 1E, simulated PSFs are shown for a fixed $A_{\mathrm{920,Gauss}} = 1$ and varying $A_{\mathrm{780,donut}}$ values of 0, 10, and 100. The case of $A_{\mathrm{780,donut}} = 0$ corresponds to conventional two-photon excitation. As $A_{\mathrm{780,donut}}$ increases, the PSF becomes narrower, resulting in a corresponding decrease in the FWHM. A density map of the FWHM as a function of $A_{\mathrm{780,donut}}$ and $A_{\mathrm{920,Gauss}}$ reveals that the FWHM is primarily determined by the ratio $A_{\mathrm{780,donut}}/A_{\mathrm{920,Gauss}}$, as indicated by the constant-FWHM contours aligned along diagonal lines, similar to Fig. 1C. This behavior arises because the apparent switching rate is set jointly by the intrinsic photoswitching rate coefficients of the rsFP and by the excitation intensities. However, unlike $k_{\mathrm{off,rsFP}}$ and $k_{\mathrm{on,rsFP}}$, the $A_{\mathrm{780,donut}}$ and $A_{\mathrm{920,Gauss}}$ contribute quadratically because the switching and excitation processes are mediated by two-photon absorption. In these simulations, $\tau_{dwell}$ was set sufficiently long, with $F_{OFF,920}/F_{ON,920}$ set to 0 and $k_{off,rsFP}$ and $k_{on,rsFP}$ set to 1 (Supplementary Table 1).

In Fig. 1F, the upper panel shows the simulated PSFs for different $\tau_{dwell}$ values ($10^{-4}$, 0.3, and 100). Increasing $\tau_{dwell}$ reduced the FWHM. At short $\tau_{dwell}$ values ($10^{-4}$, 0.3), the PSF peak is shifted along the scan direction, likely because the beam is scanned during the sequential four-photon process consisting of 2P ON-s and subsequent two-photon fluorescence excitation. Increasing $\tau_{dwell}$ further reduced the FWHM, although the achievable minimum FWHM remains unchanged because the $A_{780,donut}/A_{920,Gauss}$ ratio is kept constant at 10. However, higher $A_{780,donut}$ and $A_{920,Gauss}$ values reduce the required $\tau_{dwell}$ to achieve the improved resolution (Fig. 1F, middle). Consistently, the FWHM density map plotted as a function of $A_{780,donut}$ (at a constant intensity ratio $A_{780,donut}/A_{920,Gauss}$) and $\tau_{dwell}$ shows constant-resolution contours along diagonal lines with negative slopes, indicating that higher beam intensities compensate for shorter $\tau_{dwell}$.

**Supplementary Note 3. Membrane protrusions of MIN6 cells in pseudoislets**

Pancreatic β cells play a central role in glucose homeostasis by secreting insulin in response to elevated blood glucose levels. Dysfunction of this process leads to diabetes, one of the most prevalent metabolic disorders worldwide. Elucidating the mechanisms underlying glucose-stimulated insulin secretion is therefore of considerable physiological and clinical importance.

Pancreatic β cells form three-dimensional multicellular clusters known as islets of Langerhans *in vivo*. This architecture is essential for normal β-cell physiology and supports coordinated insulin secretion through extensive cell–cell interactions[27,28].

In the present study, we identified previously unrecognized membrane protrusions within pseudoislets (Fig. 2I and Extended Data Fig. 5). These protrusions were morphologically diverse. Some were comparatively thick and tubular (Extended Data Fig. 5C, D). The largest protrusion observed reached 1.71 μm in diameter (Extended Data Fig. 5D, F-1). Others appeared as thin, cytoneme-like protrusions no more than 238 nm in diameter, approaching the resolution limit of the present imaging system (Extended Data Fig. 5E, F-2). Some protrusions were relatively straight, whereas others were tortuous (Extended Data Fig. 5G, H). Notably, one tubular protrusion 470 nm in diameter carried a rounded, bulb-like enlargement at its distal tip (Extended Data Fig. 5H, I). The protrusions typically extended over several tens of micrometers (Extended Data Fig. 5C and G).

Although their physiological significance remains unclear, these structures may represent an additional structural element of intercellular communication within β-cell clusters. Upon glucose stimulation, mitochondrial metabolism is activated, increasing intracellular ATP levels. The resulting closure of ATP-sensitive potassium channels depolarizes the plasma membrane, leading to the opening of voltage-gated calcium channels and a rise in intracellular $Ca^{2+}$ concentration[29]. This $Ca^{2+}$ influx triggers the exocytosis of insulin-containing secretory granules, and therefore intracellular $Ca^{2+}$ dynamics are essential for insulin secretion. It has been suggested that functional heterogeneity exists within islets[30], including leader, hub, and follower β cells that coordinate collective $Ca^{2+}$ oscillations[31,32]. However, how these functionally distinct cells are structurally connected to generate coordinated activity remains unknown. The membrane protrusions identified in this study may constitute part of the structural network that underlies such coordinated multicellular behavior, although further studies will be required to determine their molecular composition and physiological function.

**Supplementary Table 1.**
**Simulation parameters for 2P-RESOLFT**

| | rsFP intrinsic properties | | | Imaging conditions | | |
|---|---|---|---|---|---|---|
| | $k_{off,rsFP}$ (μs$^{-1}$mW$^{-2}$) | $k_{on,rsFP}$ (μs$^{-1}$mW$^{-2}$) | $F_{OFF,920}/F_{ON,920}$ | $A_{780,donut}$ (mW) | $A_{920,Gauss}$ (mW) | $\tau_{dwell}$ (μs) |
| Fig. 1C upper | 0, 1, $10^2$ | $10^{-1}$ | 0 | 1 | 1 | 10000 |
| Middle | $10^{-2}$-$10^3$ | $10^{-2}$, $10^{-1}$, 1 | 0 | 1 | 1 | 10000 |
| Below (density map) | $10^{-2}$-$10^3$ | $10^{-2}$-$10^3$ | 0 | 1 | 1 | 10000 |
| Fig. 1D upper | $10^2$ | 0.1 | 0.01, 0.3, 1 | 1 | 1 | 10000 |
| Middle | 1, $10^2$, $10^3$ | 0.1 | $10^{-3}$-$10^0$ | 1 | 1 | 10000 |
| Below (density map) | $10^{-2}$-$10^3$ | 0.1 | $10^{-3}$-$10^0$ | 1 | 1 | 10000 |
| Fig. 1E upper | 1 | 1 | 0 | 0, 10, $10^2$ | 1 | 10000 |
| Middle | 1 | 1 | 0 | $10^{-2}$-$10^3$ | 0.01, 0.1, 1 | 10000 |
| Below (density map) | 1 | 1 | 0 | $10^{-2}$-$10^3$ | $10^{-2}$-$10^3$ | 10000 |
| Fig. 1F upper | 1 | 1 | 0 | 1 | 0.1* $A_{780,donut}$ | $10^{-4}$, 0.3, $10^2$ |
| Middle | 1 | 1 | 0 | $10^{-1}$, 1, 10 | 0.1* $A_{780,donut}$ | $10^{-4}$-$10^3$ |
| Below (density map) | 1 | 1 | 0 | $10^{-1}$-$10^2$ | 0.1* $A_{780,donut}$ | $10^{-4}$-$10^3$ |
| Fig. 2F | 0.74 | 1 | 1/14.1 | 0-10 | 1 | 10000 |
| Fig. 2G | $5.0 \times 10^{-5}$ | $6.8 \times 10^{-5}$ | 1/14.1 | 4.6 | 0.96 | $10^{-4}$-$10^2$ |
| | $5.0 \times 10^{-5}$ | $6.8 \times 10^{-5}$ | 1/14.1 | 9.2 | 2.1 | $10^{-4}$-$10^2$ |
| Extended Data Fig. 2 | $10^{-1}$-$10^2$ | 0.1* $k_{off,rsFP}$ | 0 | 1 | 1 | $10^{-4}$-$10^3$ |
| Extended Data Fig. 3C | 0.74 | 1 | 1/14.1 | 9.4 | 2.1 | 10000 |

| parameter | |
|---|---|
| Pixel pitch (μm) | 0.001 |
| Scan length (μm) | 5 |
| $FWHM_{920}$ (μm) | 0.452 |
| $r_{ring}$ (μm) | 0.297 |

| $P_{ON}(0)$ | 0 |
|---|---|

**Supplementary Table 2.**
**Observation conditions for 2P-RESOLFT**

| Image | 780 nm Intensity[a] (mW) | 920 nm intensity (mW) | Pixel pitch (nm) | Pixel dwell time (µs) |
|---|---|---|---|---|
| Fig. 2E | 9.4 | 2.1 | 54.44 | 1500 |
| Fig. 2F | 0 | 1.0 | 50.79 | 10000 |
| | 1.1 | 1.0 | 50.79 | 10000 |
| | 2.0 | 1.0 | 50.79 | 10000 |
| | 3.1 | 1.0 | 50.79 | 10000 |
| | 4.4 | 1.0 | 50.79 | 10000 |
| | 6.8 | 1.0 | 50.79 | 10000 |
| | 8.7 | 1.0 | 50.79 | 10000 |
| Fig. 2G (red dots) | 4.6 | 0.96 | 50.79 | 250 |
| | 4.6 | 0.96 | 50.79 | 500 |
| | 4.6 | 0.96 | 50.79 | 750 |
| | 4.6 | 0.96 | 50.79 | 1000 |
| | 4.6 | 0.96 | 50.79 | 1750 |
| Fig. 2G (Blue dots) | 9.2 | 2.1 | 50.79 | 70 |
| | 9.2 | 2.1 | 50.79 | 100 |
| | 9.2 | 2.1 | 50.79 | 175 |
| | 9.2 | 2.1 | 50.79 | 250 |
| | 9.2 | 2.1 | 50.79 | 375 |
| | 9.2 | 2.1 | 50.79 | 750 |
| Fig. 2I | 9.4 | 2.1 | 71.68 | 2000 |
| Extended Data Fig. 4 | 4.4 | 1.0 | 50.79 | 10000 |
| Extended Data Fig. 5A, middle and right | 9.4 | 2.1 | 60.4 | 2000 |
| Extended Data Fig. 5D | 9.4 | 2.1 | 75.19 | 2000 |

| | | | | |
|---|---|---|---|---|
| Extended Data Fig. 5H | 9.4 | 2.1 | 68.49 | 2000 |

[a]For conventional 2P imaging, the intensity was set to zero.
